# Association of the invasive *Haemaphysalis longicornis* tick with vertebrate hosts, other native tick vectors, and tick-borne pathogens in New York City

**DOI:** 10.1101/2020.07.01.182626

**Authors:** Danielle M. Tufts, Laura B. Goodman, Meghan C. Benedict, April D. Davis, Meredith C. VanAcker, Maria Diuk-Wasser

## Abstract

*Haemaphysalis longicornis*, the Asian longhorned tick, is an invasive ixodid tick that has rapidly spread across the northeastern and southeastern regions of the United States since first reported in 2017. The emergence of *H. longicornis* presents a potential threat for livestock, wildlife, and human health as the host associations and vector potential of this invasive pest in the United States are poorly understood. Previous field data from the United States has shown that *H. longicornis* was not associated with natural populations of small mammals or birds, but they show a preference for medium sized mammals in laboratory experiments. Therefore, medium and large sized mammals were sampled on Staten Island, New York to determine *H. longicornis* host associations and vector potential for a range of human and veterinary pathogens. A total of 97 hosts were sampled and five species of tick (*Amblyomma americanum, Dermacentor variabilis, H. longicornis, Ixodes scapularis, Ixodes cookei*) were found feeding concurrently on these hosts. *Haemaphysalis longicornis* was found in the highest proportions compared to other native tick species on raccoons (55.4%), Virginia opossums (28.9%), and white-tailed deer (11.5%). Tissue, blood, and engorged larvae were tested for 17 different pathogens using a nanoscale PCR platform. Infection with five pathogens (*Borrelia burgdorferi, Anaplasma phagocytophilum, Rickettsia* spp., *Mycoplasma haemocanis*, and *Bartonella* spp.) was detected in host samples, but no pathogens were found in any larval samples. These results suggest that although large and medium sized mammals feed large numbers of *H. longicornis* ticks in the environment there is presently a low potential for *H. longicornis* to acquire pathogens from these wildlife hosts.

**Highlights:**

- *H. longicornis* were sampled from seven genera of large and medium-sized mammals
- Raccoons, opossums, and white-tailed deer fed a large proportion of *H. longicornis*
- *H. longicornis* did not acquire pathogens through co-feeding with native tick vectors
- Host species were infected with a range of pathogens of human and veterinary concern
- Host-derived *H. longicornis* engorged larvae were not infected with any pathogens

## 1. Introduction

The invasive Asian longhorned tick, *Haemaphysalis longicornis*, has become established in the northeastern and southeastern parts of the United States (Beard et al., 2018; Hutcheson et al., 2019). This invasive pest has spread quickly across 12 states in recent years, most likely because of its ability to reproduce parthenogenetically and feed on a wide range of hosts (Chen et al., 2012; Heath, 2016; Rainey et al., 2018; Beard et al., 2018). In its native range and other previously introduced regions (Australia, New Zealand, Pacific Islands), *H. longicornis* has been recovered from a wide range of wildlife hosts (including mice, rats, hares, possums, deer, and birds) and livestock hosts (cattle, sheep, goats) as well as humans (Heath, 2016). However, in a field study conducted on Staten Island, New York (NY) from 2017-2018, no *H. longicornis* of any life stage were found feeding on seven different species of bird or on any small rodent species (white-footed mice, brown rats, eastern chipmunks, and northern short-tailed shrews) despite high abundances of all three life stages questing in the same environments and feeding on white-tailed deer (Tufts et al., 2019). Tufts and colleagues (2019) suggested a low likelihood that *H. longicornis* would acquire regional rodent-associated pathogens of human concern due to the absence of feeding on the dominant reservoir hosts for these pathogens (including *Peromyscus leucopus* mice). In a recent laboratory study, *H. longicornis* showed a significant preference towards domestic cat, dog, and white-tailed deer hair while avoiding *P. leucopus* and human hair (Ronai et al., 2020). In the United States, the Asian longhorned tick has been associated with a small number of domestic livestock hosts (sheep, goats, cattle, horses), domestic pets (cats and dogs), and a small number of wildlife hosts (white-tailed deer, raccoon, Virginia opossum, carnivores, Eastern cottontail; Tufts et al., 2019; USDA, 2020). Additionally, a recent report suggested *H. longicornis* killed, via exsanguination, five cattle in North Carolina (Neault, 2019). These studies suggest that the United States populations of *H. longicornis* may preferentially feed on large or medium sized mammalian wildlife and domestic companion or livestock hosts.

Many native large and medium sized mammalian hosts (i.e. eastern cottontail rabbit, feral cats, gray squirrels, raccoons, striped skunks, Virginia opossum, white-tailed deer) can serve as reservoirs for pathogens of human and veterinary concern (Fish and Dowler, 1989; Levin et al., 2002; LoGiudice et al., 2003) and might constitute an alternative route for the acquisition of these pathogens in *H. longicornis*. White-tailed deer (*Odocoileus virginianus*) are considered reservoir hosts of *Ehrlichia chaffeensis*, the causative agent of human monocytic ehrlichiosis, *Ehrlichia ewingii*, the causative agent of granulocytic ehrlichiosis in humans and canines, *Borrelia miyamotoi*, a spirochete responsible for relapsing fever, *Anaplasma* spp., and *Babesia* spp. pathogens (Lockhart et al., 1997; Comer et al., 2001; Yabsley et al., 2002; Massung et al., 2005; Inokuma, 2007; Han et al., 2016). A large number of *H. longicornis* were observed feeding on white-tailed deer individuals in New York City (Tufts et al., 2019) which could lead to the transmission of the aforementioned pathogens to this newly invaded tick. Although, white-tailed deer are not competent hosts for *Borrelia burgdorferi*, a causative agent of Lyme disease (Telford et al., 1988; Pfaffle et al., 2013), *H. longicornis* may acquire these and other pathogens via co-feeding spatiotemporally with infected native tick species.

While evidence is mounting that the invasive Asian longhorned tick has the ability to feed on a wide range of hosts in the United States, its potential to serve as a vector for pathogens of human and veterinary concern in the United States is still poorly understood. In its native range, *H. longicornis* is known to vector or has been found associated with several pathogens (i.e. severe fever with thrombocytopenia syndrome virus, *Anaplasma* spp., *Rickettsia* spp., *Babesia* spp., *Theileria* spp., *Borrelia* spp., etc.), including pathogens in the same genus and species as some endemic pathogens in the United States (Chu et al., 2008; Luo et al., 2015; Heath, 2016; Zhuang et al., 2018; Eisen, 2020; Zheng et al., 2020). Recent evidence from a laboratory study on *Mus musculus* mice illustrated that *H. longicornis* can acquire *B. burgdorferi* sensu stricto (s.s.) spirochetes in the larval life stage; however, the bacteria were unable to persist through molting into the nymphal life stage (Breuner et al., 2020), suggesting that *H. longicornis* in the United States is unable to vector this pathogen. Another recent laboratory study found that larval and nymphal *H. longicornis* were able to acquire *Rickettsia rickettsii*, the causative agent of Rocky Mountain spotted fever, from infected guinea pigs and transmit the pathogen to susceptible guinea pigs; they also observed transovarial transmission of *R. rickettsii* at a low frequency (Stanley et al., 2020). Although capable of serving as experimental vectors of *R. rickettsii*, the ability of natural populations of *H. longicornis* to vector this pathogen is still undetermined as no naturally infected *H. longicornis* ticks have been documented in the United States. While the role of *H. longicornis* as a vector of human pathogens is uncertain, there is increasing evidence of its potential veterinary importance. Infection with *Theileria orientalis* Ikeda genotype, an important pathogen of cattle, was found in 13% of questing *H. longicornis* nymphs in Virginia where a cattle herd was recently affected by theileriosis (Thompson et al., 2020), further supporting the feeding habits of *H. longicornis* for larger mammalian hosts and providing insights into their vector potential.

To better understand the host associations and potential to acquire pathogens directly from hosts or through co-feeding of the invasive Asian longhorned tick in the New York City area a field study was conducted to i) assess which large and mesomammalian wildlife hosts *H. longicornis* of all life stages feeds on, ii) assess which other tick species are co-feeding spatiotemporally with *H. longicornis*, and iii) investigate the pathogen prevalence of hosts and host-derived *H. longicornis* engorged larvae for local pathogens of human and veterinary health concern. This information will greatly increase our understanding of the type of health threat *H. longicornis* poses to wildlife, livestock, domestic pets, and human hosts.

## 2. Materials and Methods

### 2.1 Sample collection

Mesomammals were trapped in six city parks throughout Staten Island, NY (Clay Pit Ponds State Park, Conference House Park, Great Kills Park, Latourette Park, Mount Loretto Unique Area, and Willowbrook Park) from June-September 2019 during bi-weekly, two night trapping sessions. A maximum of two parks were visited during a trapping session and parks were sampled 1-3 times during the trapping period (Willowbrook Park sampled 26 Jun; Conference House Park sampled 16 Jul, 13 Aug, and 9 Sep; Clay Pit Ponds Park sampled 30 Jul and 9 Sep; Great Kills Park sampled only on 26 Aug; Latourette Park sampled 26 Aug and 17 Sep; Mount Loretto Unique Area sampled 17 Sep). Tomahawk live traps (30.5H x 33.0W x 81.3L cm) were set at dusk and checked at dawn; traps were baited with either wet cat food or peanut butter. Mesomammals were anesthetized in the field using Telazol (dosage adjusted for species and individual body weight). While under anesthesia, sections of the body were examined separately (right ear, left ear, head/body), ticks were removed and stored in 100% ethanol in separate microcentrifuge tubes from each body section. Tick collection continued until all visible ticks were collected, however some ticks may have inadvertently been missed. A small ear biopsy was collected from the pinna and stored in 100% ethanol, each animal was given a unique tag number, and whole blood (no preservation media), serum (collected using BD Microtainer Serum Separator Tubes, Fisher Scientific, Waltham, MA, USA), fecal, and saliva samples were collected from each animal. Once all samples were collected, animals were placed back in the traps and allowed to fully recover from anesthesia before being released at the site of capture.

As previously described (Tufts et al., 2019), white-tailed deer (*Odocoileus virginianus*) were opportunistically sampled in August 2018 as part of a sterilization project aimed for population reduction. Trapping locations of white-tailed deer were from four locations on Staten Island, NY (Clay Pit Ponds Park, the College of Staten Island, Freshkills Park, and Mount Loretto Unique Area). Once the animals were anesthetized, attached ticks were removed from the antlers, right ear, left ear, head, and body. Ticks were the only samples collected from these individuals.

Fecal samples were collected by rectal swab of the anesthetized mesomammals and preserved in a 0.9% potassium dichromate solution. A fecal float was performed using Sheather’s solution and centrifuging samples at 1500 rpm for 10 min. Samples were immediately viewed using a light microscope (Revolve, Echo Laboratories, San Diego, CA) at 40x magnification and pictures were taken for morphological identification of macroparasites.

Saliva samples were collected by rubbing a sterile cotton applicator in the mouth of the anesthetized mesomammals and placing the swab in a tube containing 1mL of Eagles Growth Media (Appler et al., 2019). These samples were screened at the New York State Department of Health, Rabies Laboratory for rabies and canine distemper using a previously designed real-time PCR method (Elia et al., 2006; Appler et al., 2019).

All procedures and protocols were approved by the Columbia University Institutional Animal Care and Use Committee (IACUC) and all New York State and City permits were obtained prior to completion of this work.

### 2.2 Mesomammal and tick DNA extraction and pathogen screening

Ear biopsy tissues and *H. longicornis* engorged larvae were processed individually following a previously described protocol (Yuan et al., in press). Briefly, samples were homogenized in 400 µl of phosphate-buffered saline (PBS) at pH 7.4 (Gibco, Gaithersburg, MD, USA) along with a 4 mm hollow brass bead (Hareline Dubbin, Monroe, OR) and homogenized for 5 min at 2,100 rpm in a Mini-Beadbeater-96 (BioSpec Products, Bartlesville, OK, USA). A 175 µl aliquot of each sample was extracted using MagMAX total nucleic acid isolation (Applied Biosystems, Foster City, CA, USA) on an automated extraction Kingfisher Flex instrument (ThermoFisher Scientific, Waltham, MA, USA). For *H. longicornis* engorged larvae samples, DNA extraction aliquots from 10 ticks were pooled together and up to 20 larvae were analyzed per individual host.

DNA was extracted from whole blood samples using a custom MagMAX with clarifying solution kit (Applied Biosystems, Foster City, CA, USA), made by combining the MagMAX CORE nucleic acid purification kit with the MagMAX CORE mechanical lysis module clarifying solution in place of the lysis buffer provided in the kit. Blood samples were first chemically lysed by adding 235 µl of clarifying solution and 10 µl of proteinase K to each sample with a total volume less than 300 µl of blood and 352.5 µl of clarifying solution and 15 µl of proteinase K to samples with a total volume greater than or equal to 300 µl. Carrier RNA (ThermoFisher Scientific, Waltham, MA, USA), bacteriophage MS2 (an internal positive control), and Qiagen DX reagent (Qiagen, Germantown, MD, USA) were added to each tube. Samples were then vortexed and incubated at 56°C for 18 h. After incubation, samples were vortexed and DNA was extracted using the aforementioned protocol.

For all samples (tissue, whole blood, and *H. longicornis* engorged larvae) two negative controls (PBS buffer only) were included with each extraction plate. Samples were screened for 17 pathogens of human and veterinary health concern (*Anaplasma phagocytophilum, Anaplasma marginale, Babesia microti, Bartonella spp*., *B. burgdorferi* s.s., *Borrelia mayonii, B. miyamotoi, Ehrlichia canis, E. chaffeensis, E. ewingii, Mycoplasma haemocanis*, Powassan virus, *Rickettsia spp, R. rickettsii*, Severe Fever with Thrombocytopenia Syndrome Virus (SFTSV), *T. orientalis*, and Heartland virus) using an OpenArray Tick Nanochip workflow described elsewhere (Goodman et al., 2016).

### 2.3 Statistical Analyses

A one-way analysis of variance (ANOVA) was used to assess differences in the total numbers of ticks of the three most abundant tick species collected from the most abundant host species. A post-hoc Tukey’s honestly significant difference (HSD) test was incorporated to determine pairwise differences between specific tick species.

## 3. Results

### 3.1 Mammal trapping and ectoparasite burden

A total of 97 animals were sampled from Staten Island, NY belonging to seven different mammalian species (domestic cat – *Felis catus*; Eastern gray squirrel – *Sciurus carolinensis*; marmot – *Marmota monax*; raccoon – *Procyon lotor*; striped skunk – *Mephitis mephitis*; Virginia opossum – *Didelphis virginiana*; white-tailed deer – *O. virginianus*), with raccoons accounting for the majority of individuals (Table 1). Mesomammals were sampled in different proportions from each of the six park locations (Fig. 1). Ectoparasites collected from all animals consisted of ticks (*n* = 12,081), fleas (*n* = 60), and lice (*n* = 12). Five species of tick were collected from all hosts and *H. longicornis* accounted for the highest proportion of all ticks collected (*n* = 7,442/12,081; 61.6%) followed by *A. americanum* (*n* = 3,088/12,081; 25.6%), *I. scapularis* (*n* = 1,510/12,081; 12.5%), *Ixodes cookei* (*n* = 32/12,081; 0.26%), and *Dermacentor variabilis* (*n* = 4/12,081; 0.03%; Table 1). Because *I. cookei* and *D. variabilis* were observed infrequently they were not included in further analyses. The proportions of the remaining tick species varied depending on host species; the infestation intensity of *I. scapularis* was highest on white-tailed deer, while the infestation intensity of *A. americanum* and *H. longicornis* was highest in raccoons (Table 2; Fig. 2). Focusing on the three host species that fed the highest number of ticks of any life stage at all locations, significantly more *H. longicornis* (*n* = 4,126) were recovered from all raccoons compared to *I. scapularis* (*n* = 121; *P* < 0.01, degrees of freedom (df) = 116, *F* = 6.34), but not compared to *A. americanum* (*n* = 2,749), significantly more *H. longicornis* (*n* = 2,152) were collected from all opossums compared to *A. americanum* (*n* = 70; *P* < 0.01, df = 62, *F* = 5.97) while no difference was observed for *I. scapularis* (*n* = 797), and significantly more *H. longicornis* (*n* = 855) were removed from all white-tailed deer compared to *A. americanum* (*n* = 108; *P* < 0.01, df = 47, *F* = 5.58), but not in comparison to *I. scapularis* (*n* = 550; Fig. 2). The proportion and distribution of these three tick species also varied geographically (Fig. 3); however statistical analyses could not be performed based on the collected data. Within host species groups, *H. longicornis* was found co-feeding on the same host body section with one of the other tick species (*I. scapularis*: IS or *A. americanum*: AA) on feral cats (*n*_IS_ = 3, 60.0%; *n*_AA_ = 4, 80.0%), marmots (*n*_IS_ = 1, 50.0%; *n*_AA_ = 2, 100.0%), opossums (*n*_IS_ = 20, 95.2%; *n*_AA_ = 8, 38.1%), raccoons (*n*_IS_ = 18, 46.2%; *n*_AA_ = 34, 87.2%), striped skunks (*n*_IS_ = 1, 50.0%; *n*_AA_ = 1, 50.0%), and white-tailed deer (*n*_IS_ = 13, 81.3%; *n*_AA_ = 10, 62.5%) (Suppl Table 1).

**Table 1.** Number of host individuals sampled and the number of each tick life stage and species collected from all sampled hosts on Staten Island, NY (2018-2019).

| Host species | *n* | *I. scapularis* |  |  | *A. americanum* |  |  | *H. longicornis* |  |  | Total |
| --- | --- | --- | --- | --- | --- | --- | --- | --- | --- | --- | --- |
|  |  | A | N | L | A | N | L | A | N | L |  |
| Cat | 5 | 0 | 1 | 3 | 0 | 1 | 37 | 0 | 3 | 247 | 292 |
| Gray Squirrel | 12 | 0 | 5 | 14 | 0 | 0 | 0 | 0 | 0 | 2 | 21 |
| Marmot | 2 | 0 | 8 | 1 | 0 | 5 | 25 | 0 | 0 | 30 | 69 |
| Opossum | 21 | 0 | 28 | 769 | 1 | 5 | 64 | 58 | 534 | 1560 | 3019 |
| Raccoon | 39 | 0 | 19 | 102 | 2 | 244 | 2503 | 232 | 133 | 3761 | 6996 |
| Striped skunk | 2 | 0 | 1 | 9 | 1 | 10 | 82 | 0 | 0 | 27 | 130 |
| White-tailed deer* | 16 | 12 | 6 | 532 | 12 | 13 | 83 | 168 | 74 | 613 | 1513 |
| Total | 97 | 12 | 68 | 1430 | 16 | 278 | 2794 | 458 | 744 | 6240 | 12040 |

A = adult, N = nymph, L = larvae; \*White-tailed deer were sampled only in 2018

**Table 2.**
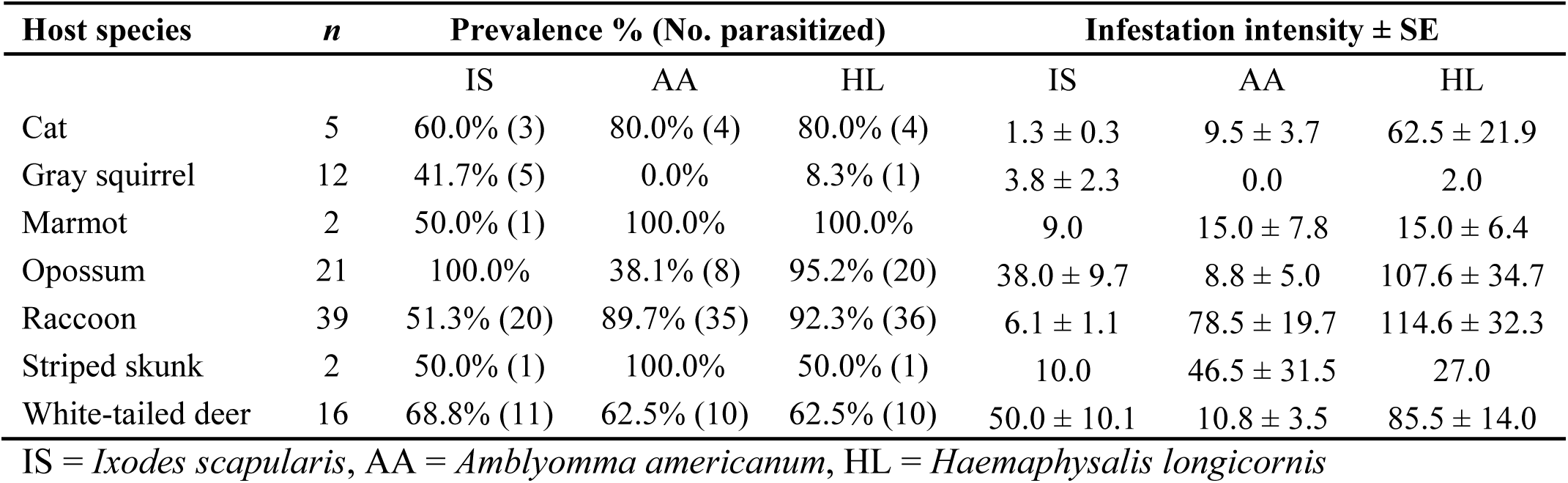
Infestation prevalence (number of infested individuals/total individuals sampled) along with the number of parasitized host individuals by a particular tick species (shown in parentheses) and infestation intensity (mean ± standard error of the number of ticks per host) for each species of tick collected from each host species.

**Figure 1.**
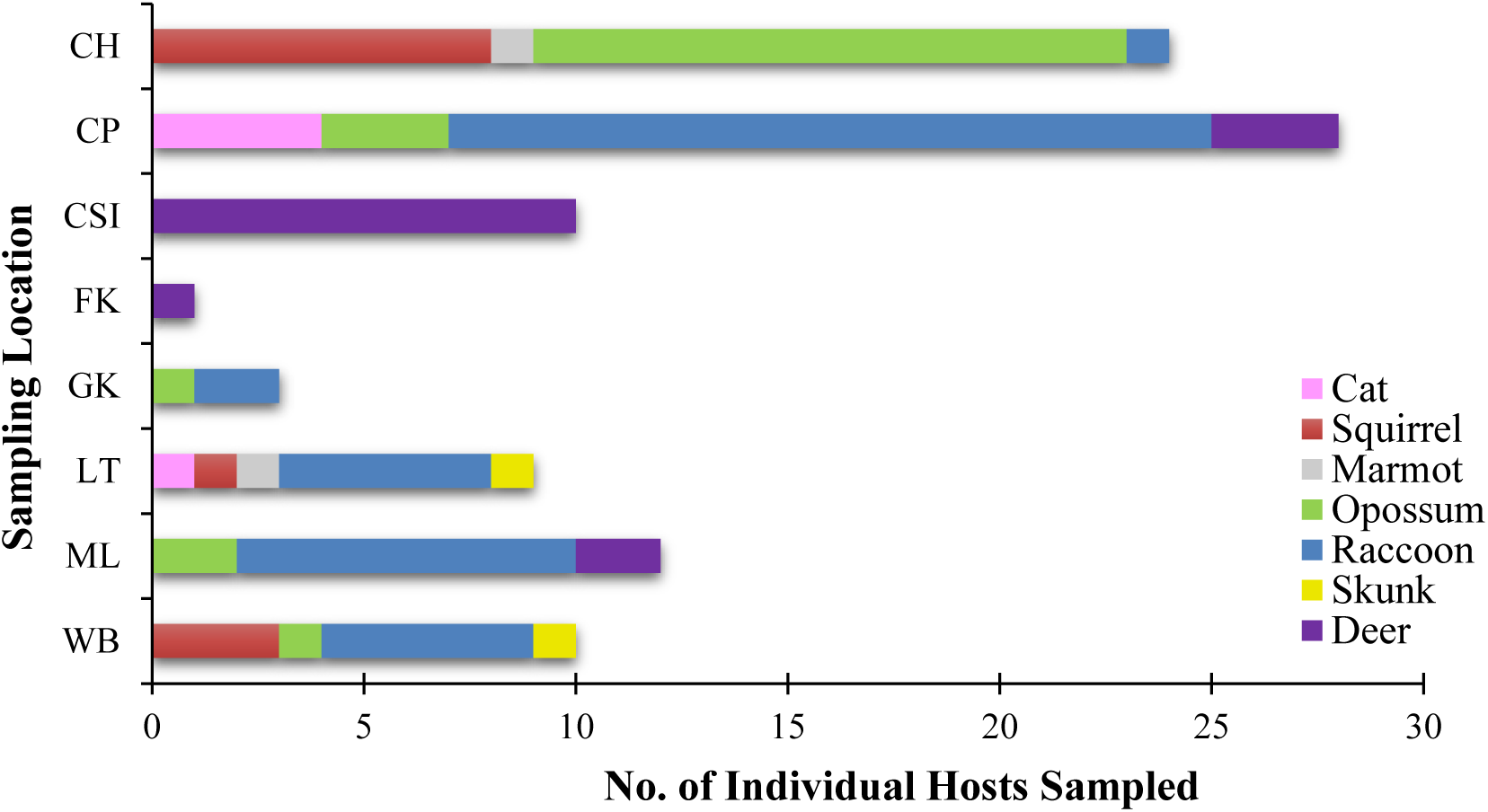
Number of individuals from each host species sampled at each park location on Staten Island, NY (2018-2019).

**Figure 2.**
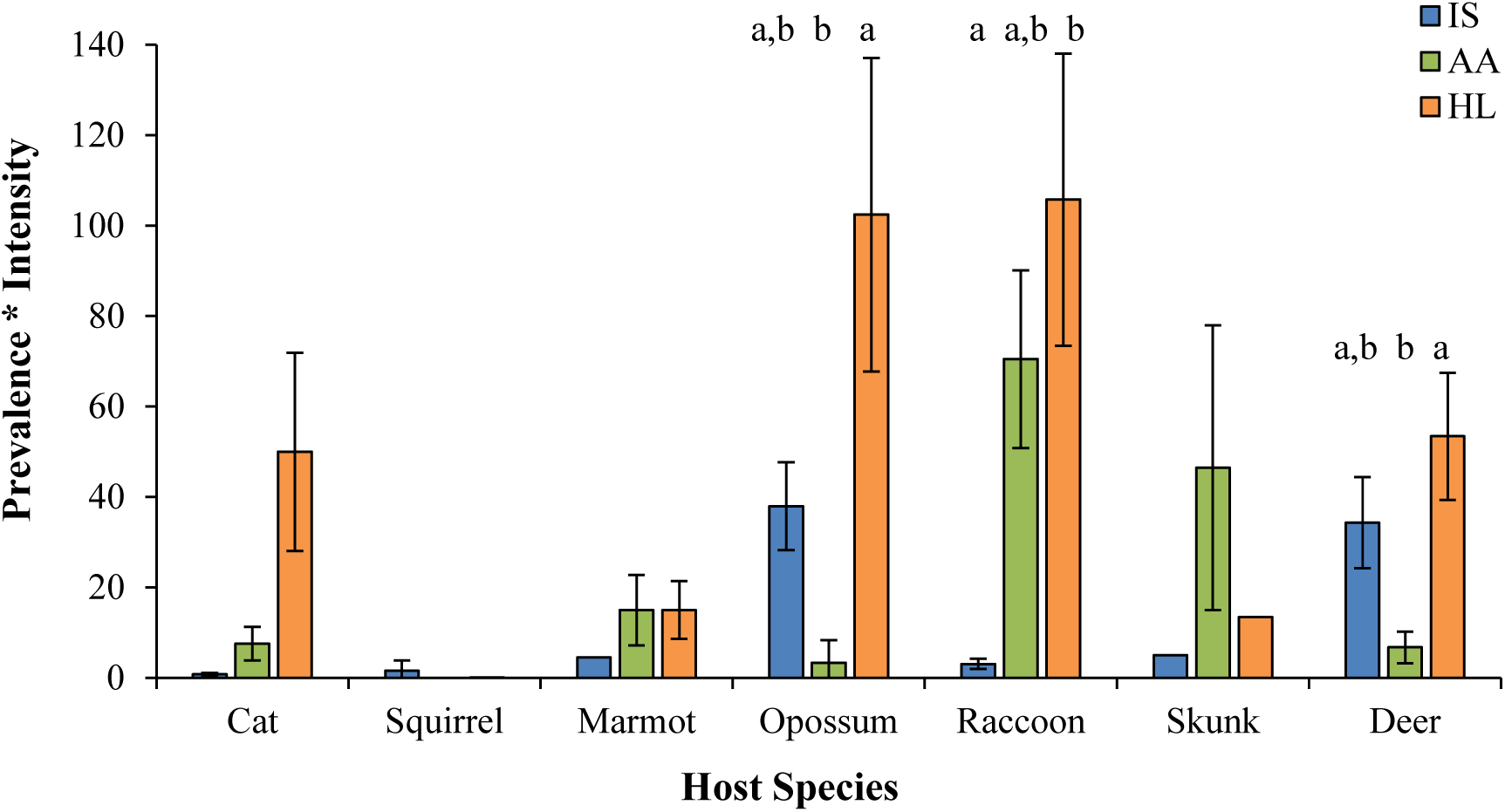
Infestation intensity of the three major ticks species (IS = *Ixodes scapularis*, AA = *Amblyomma americanum*, HL = *Haemaphysalis longicornis*) collected from each host species sampled on Staten Island. Different letters denote significant differences (*P* < 0.01) between the numbers of ticks collected from each host species.

**Figure 3.**
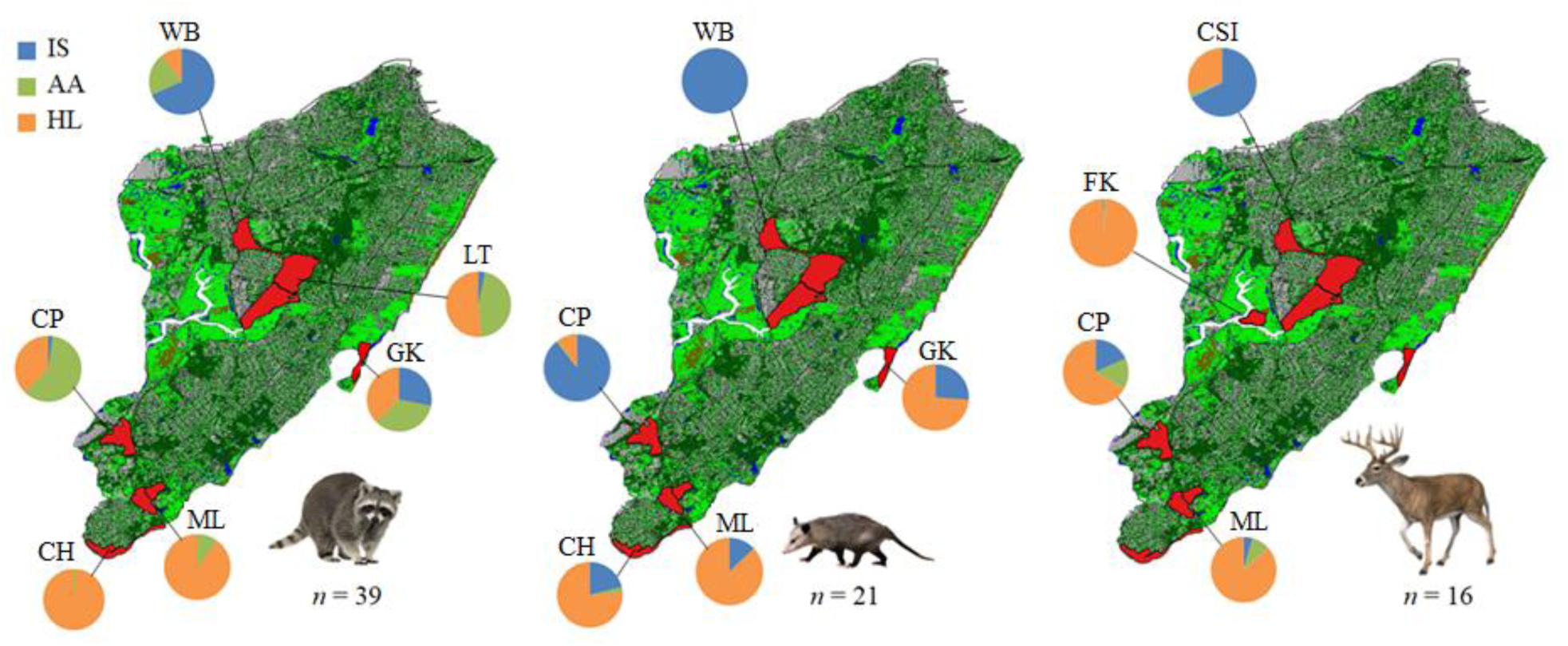
Proportion of individual ticks species (IS = *Ixodes scapularis*, AA = *Amblyomma americanum*, HL = *Haemaphysalis longicornis*) over all ticks collected from the most heavily infested host species (raccoons, opossums, and white-tailed deer) at each trapping location on Staten Island (WB = Willowbrook Park, CSI = College of Staten Island, LT = Latourette Park, FK = Freshkills Park, GK = Great Kills Park CP = Clay Pit Ponds State Park, ML = Mount Loretto Unique Area, CH = Conference House Park).

### 3.2 Mesomammal pathogen screening

Infection screening from ear biopsy tissue samples (*n* = 77) revealed that 23.4% of individuals were infected with at least one pathogen. Six individuals were infected with *A. phagocytophilum* (7.8% infection prevalence; *n* = 5 raccoons, *n* = 1 marmot), two individuals were infected with *B. burgdorferi* (2.6% prevalence; *n* = 1 raccoon, *n* = 1 skunk), nine raccoons were infected with *M. haemocanis* (11.7% prevalence), and three raccoons were infected with a *Rickettsia* spp. (3.9% prevalence; Table 3).

**Table 3.** Infestation prevalence (number of infested individuals/total individuals and total number of infested hosts in parentheses) and infestation intensity (mean ± standard error of the number of ticks per host) for hosts and tick species collected at each park location (italicized heading).

| Host Species | *n* | Prevalence (No. parasitized) |  |  | Infestation intensity ± SE |  |  |
| --- | --- | --- | --- | --- | --- | --- | --- |
|  |  | IS | AA | HL | IS | AA | HL |
| *Willowbrook Park* |  |  |  |  |  |  |  |
| Squirrel | 3 | 33.3% (1) | 0.0% | 0.0% | 1.0 | 0.0 | 0.0 |
| Opossum | 1 | 100.0% | 0.0% | 0.0% | 3.0 | 0.0 | 0.0 |
| Raccoon | 5 | 80.0% (4) | 60.0% (3) | 40.0% (2) | 3.3 ± 1.2 | 1.3 ± 0.3 | 1.0 |
| Skunk | 1 | 0.0% | 100.0% | 0.0% | 0.0 | 2.0 | 0.0 |
| *College of Staten Island* |  |  |  |  |  |  |  |
| White-tailed Deer | 10 | 100.0% | 40.0% (4) | 80.0% (8) | 44.9 ± 13.6 | 3.5 ± 1.6 | 25.1 ± 9.9 |
| *Latourette Park* |  |  |  |  |  |  |  |
| Cat | 1 | 100.0% | 100.0% | 100.0% | 1.0 | 3.0 | 29.0 |
| Squirrel | 1 | 100.0% | 0.0% | 0.0% | 1.0 | 0.0 | 0.0 |
| Marmot | 1 | 100.0% | 100.0% | 100.0% | 9.0 | 26.0 | 6.0 |
| Raccoon | 5 | 40.0% (2) | 100.0% | 100.0% | 7.5 ± 2.5 | 40.8 ± 18.7 | 46.4 ± 34.3 |
| Skunk | 1 | 100.0% | 100.0% | 100.0% | 10.0 | 91.0 | 27.0 |
| *Freshkills Park* |  |  |  |  |  |  |  |
| White-tailed Deer | 1 | 100.0% | 100.0% | 100.0% | 1.0 | 1.0 | 161.0 |
| *Great Kills Park* |  |  |  |  |  |  |  |
| Opossum | 1 | 100.0% | 0.0% | 100.0% | 18.0 | 0.0 | 51.0 |
| Raccoon | 2 | 100.0% | 100.0% | 100.0% | 4.5 ± 0.4 | 5.5 ± 1.8 | 6.0 ± 2.8 |
| *Clay Pit Ponds Park* |  |  |  |  |  |  |  |
| Cat | 4 | 50.0% (2) | 75.0% (3) | 75.0% (3) | 1.5 ± 0.4 | 11.7 ± 4.3 | 73.7 ± 26.2 |
| Opossum | 3 | 100.0% | 33.3% (1) | 100.0% | 69.7 ± 52.0 | 2.0 | 7.7 ± 3.8 |
| Raccoon | 18 | 66.7% (12) | 100.0% | 100.0% | 7.0 ± 1.7 | 125.7 ± 34.6 | 79.1 ± 22.8 |
| White-tailed Deer | 3 | 100.0% | 100.0% | 100.0% | 30.7 ± 13.2 | 25.7 ± 4.5 | 112.0 ± 20.0 |
| *Mt Loretto Unique Area* |  |  |  |  |  |  |  |
| Opossum | 2 | 100.0% | 0.0% | 100.0% | 5.0 ± 2.1 | 0.0 | 33.5 ± 10.3 |
| Raccoon | 8 | 0.0% | 87.5% (7) | 100.0% | 0.0 | 37.9 ± 15.3 | 288.5 ± 113.1 |
| White-tailed Deer | 2 | 50.0% | 100.0% | 100.0% | 8.0 | 8.0 ± 1.4 | 78.5 ± 1.8 |
| *Conference House Park* |  |  |  |  |  |  |  |
| Squirrel | 8 | 37.5% (3) | 0.0% | 12.5% (1) | 5.7 ± 3.4 | 0.0 | 2.0 |
| Marmot | 1 | 0.0% | 100.0% | 100.0% | 0.0 | 4.0 | 24.0 |
| Opossum | 14 | 100.0% | 50.0% (7) | 100.0% | 39.8 ± 7.2 | 9.7 ± 5.4 | 143.6 ± 46.2 |
| Raccoon | 1 | 0.0% | 100.0% | 100.0% | 0.0 | 2.0 | 148.0 |

IS = *Ixodes scapularis*, AA = *Amblyomma americanum*, HL = *Haemaphysalis longicornis*

From blood sample analysis (*n* = 64), 46.9% of individuals screened were infected with at least one pathogen. Seventeen individuals were infected with *A. phagocytophilum* (26.6% infection prevalence; *n* = 15 raccoons, *n* = 1 marmot, *n* = 1 feral cat), two individuals were infected with a *Bartonella* spp. (3.1% prevalence; *n* = 1 marmot, *n* = 1 gray squirrel), 14 raccoons were infected with *M. haemocanis* (21.9% prevalence), and one raccoon was infected with a *Rickettsia* spp. (1.6% infection prevalence; Table 3). Of all pathogen infected individuals (*n* = 33), 78.8% were infected with one pathogen (*n* = 26), 18.2% were coinfected with two pathogens (*n* = 6), and 3.0% were coinfected with three pathogens (*n* = 1) (Table 4). Infection prevalence varied greatly for each species and location of capture (Fig. 4 A,B). Tissue and blood samples that were positive for a *Rickettsia* spp. were specifically screened for *R. rickettsii* (Kato et al., 2013; Yuan et al., in press) and all samples were negative. Host infection status (tissue and blood) was determined through a real-time PCR approach and may not represent an active infection in these animals.

**Table 4.**
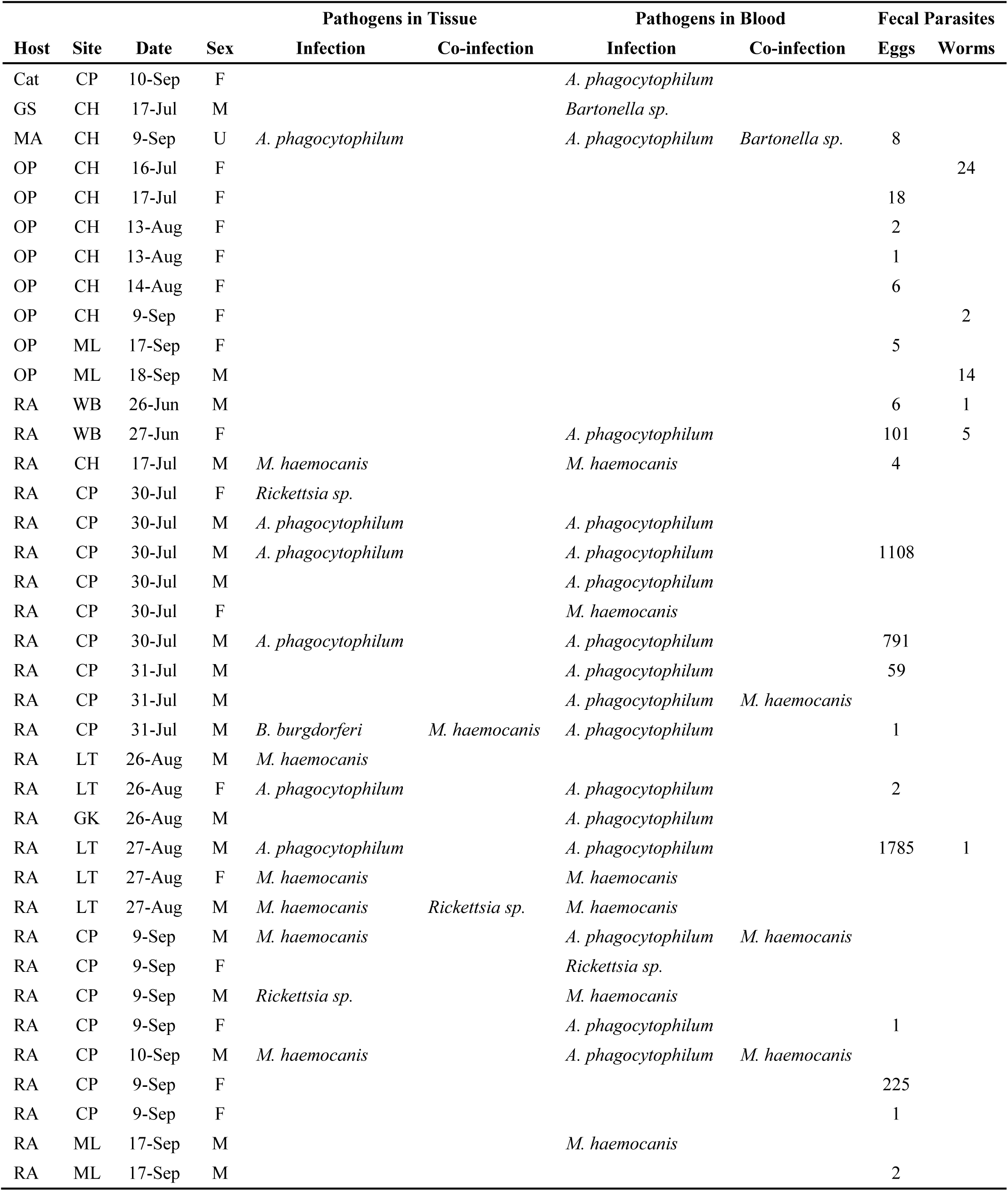
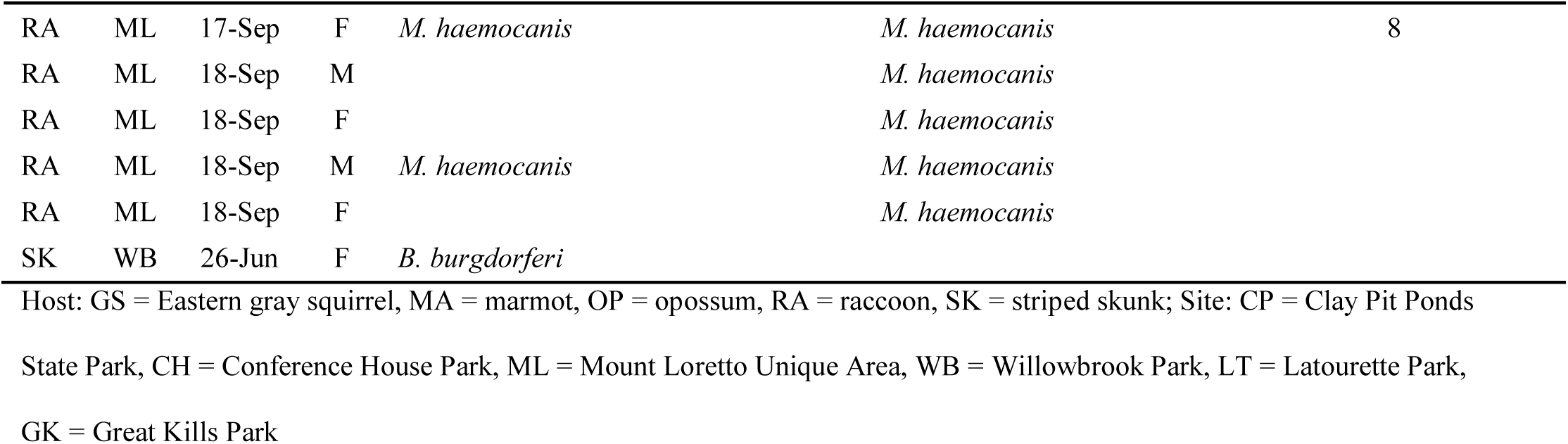
Pathogen species detected in ear biopsy tissue, whole blood, and fecal samples in all mesomammals sampled.

**Figure 4.**
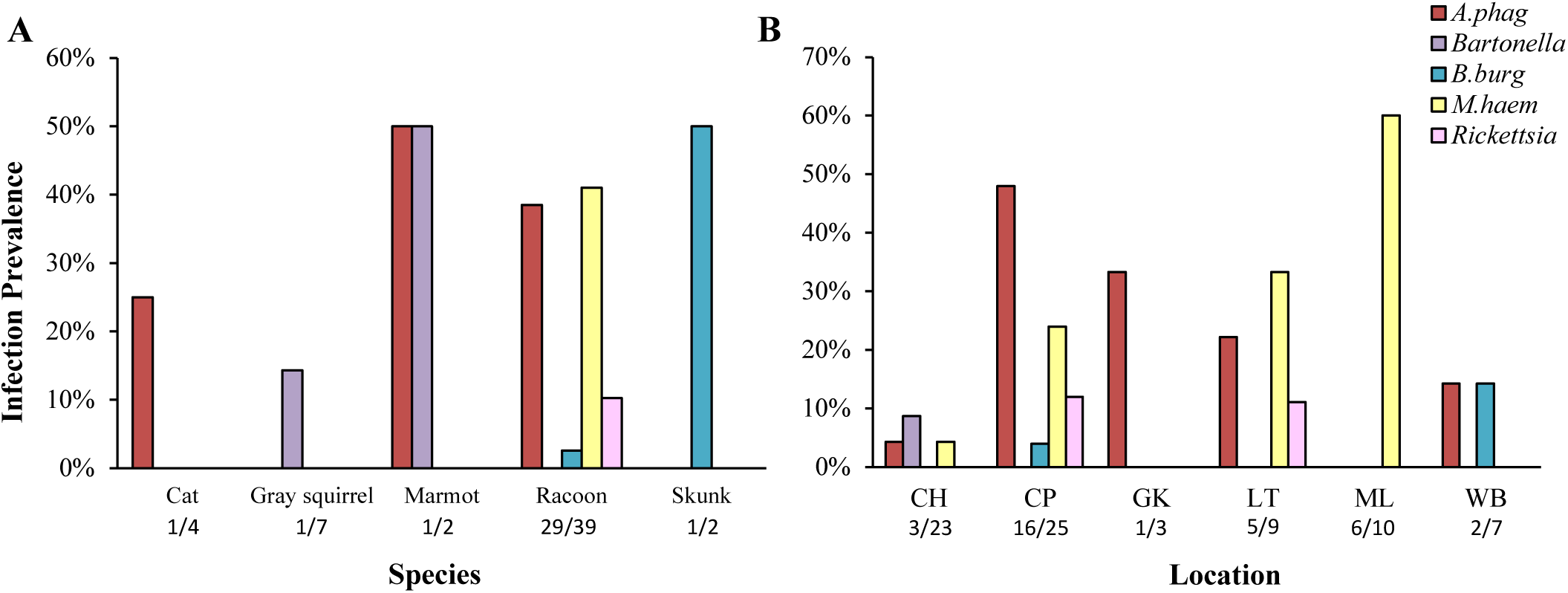
Infection prevalence of each pathogen in all individual hosts tested, sorted by (A) host species and (B) sampling location on Staten Island (CH = Conference House Park, CP = Clay Pit Ponds State Park, GK = Great Kills Park, LT = Latourette Park, ML = Mount Loretto Unique Area, WB = Willowbrook Park). Numbers on the x-axis denote the number of infected individuals/number of individuals tested.

Analysis of fecal samples (*n* = 71) revealed that 28.2% of animals screened were infected with nematode eggs of the genus *Trichuris, Toxocara, Toxascaris*, and/or *Baylisascaris* (*n* = 20 hosts), and 8.5% of screened individuals were infected with strongylid nematodes (*n* = 6 hosts; Table 4). While none of the opossums were infected with any microbial pathogens, 38.09% had fecal macroparasites (*n* = 8 hosts; Table 4).

All saliva samples (*n* = 69) tested negative for rabies and canine distemper. However, saliva is a not a definitive method of testing for rabies or distemper, and the lack of positive results does not necessarily indicate the testing location is free of either virus.

### 3.3 H. longicornis *pathogen screening*

A subset of partially and fully engorged *H. longicornis* larvae (*n* = 813), collected from both infected and uninfected hosts, tested negative for each of the 17 human and veterinary pathogens, regardless of engorgement level.

## 4. Discussion

Seven different genera of large and medium sized mammals were sampled on Staten Island, NY in 2018-2019 and five species of ixodid ticks were collected from these hosts. The most abundant species of tick removed from mammalian hosts were *I. scapularis, A. americanum*, and *H. longicornis* which were found feeding in close space and time (co-feeding) to each other throughout the trapping season. Raccoons, opossums, and white-tailed deer fed the largest proportion of *H. longicornis* ticks compared to the other host species sampled. The highest infestation prevalence and infestation intensity of *H. longicornis* was observed from raccoons. Hosts were infected with a range of pathogens; however no *H. longicornis* engorged larvae were infected with any pathogens, suggesting a limited role of the Asian longhorned tick as a significant vector of these pathogens in the United States.

The newly invaded *H. longicornis* ticks were introduced into the eastern United States presumably after 2010 (Beard et al., 2018) and potentially via three invasion events originating from east Asia on domestic pets (Egizi et al., in press). *Ixodes scapularis* and *A. americanum* ticks are indigenous to the United States and are primarily found in the eastern half of the country. The geographic distribution of these ticks and the dominant habitat and the wildlife host community on Staten Island, NY are conducive for *H. longicornis* to co-occur with these other native tick species. Concurrent feeding of *H. longicornis* with at least one of the native tick species was observed for 78% of hosts sampled, highlighting the potential for cross-tick species transmission through co-feeding of human and veterinary pathogens.

Raccoons fed significantly fewer *I. scapularis* (*n* = 121) and only immature life stages compared to all life stages of *H. longicornis* (*n* = 4,126), for example a total of 929 ticks were removed from one individual raccoon (*n* = 876 *H. longicornis, n* = 50 *A. americanum, n* = 0 *I. scapularis*). Raccoons, when infested with *I. scapularis*, may elicit an immune response to tick salivary proteins which significantly decreases the number of attached ticks (Craig et al., 1996). As *H. longicornis* are a newly invaded species, raccoons may not elicit or may show a reduced host acquired tick resistance which might explain why significantly fewer *I. scapularis* were recovered from raccoons in comparison with *H. longicornis*. Opossum and white-tailed deer fed significantly fewer *A. americanum* (*n* = 70; *n* = 108, respectively) compared to *H. longicornis* (*n* = 2,152; *n* = 855, respectively). Recovering fewer *A. americanum* from opossums is consistent with other studies (Childs and Paddock, 2003); however, over 3,000 ticks of four different species were recovered from opossums, suggesting that opossums might not be very effective tick groomers (as proposed by Keesing et al., 2010) or that certain tick species avoid feeding on opossums (i.e. *A. americanum*). Tick samples from white-tailed deer were collected only in August 2018 compared to mesomammals that were sampled from June-September in 2019 making statistical analyses between hosts impractical because of introduced biases based on seasonal tick density fluctuations and trapping effort.

*Anaplasma phagocytophilum* was the most prevalent and widespread pathogen identified from mesomammals on Staten Island and infected three different species: a feral cat (25.0%), a marmot (50.0%), and several raccoons (38.5%, *n* = 15/39) and was found infecting at least one host individual in all but one park. Similarly, Levin and colleagues (2002) investigated the reservoir potential of medium-sized mammals and found *A. phagocytophilum* infection in the blood of 33.3% of sampled feral cats and 24.6% of sampled raccoons with no infection in other sampled hosts (gray squirrels, striped skunks, and Virginia opossums). Data presented in this study suggests that feral cats and raccoons are important hosts of *A. phagocytophilum* with potential for transmission to *H. longicornis* and other tick species on Staten Island.

The second most abundant pathogen recovered from Staten Island mammalian hosts was *M. haemocanis*, a hemoplasma erythrocytic pathogen commonly found infecting immunosuppressed canines. This pathogen poses a potential hazard to domestic pets and was detected only in raccoons (41.0%, *n* = 16/39) from four parks. Limited information is available about *M. haemocanis* in raccoons in the United States; however, one study from Georgia found 62.1% of urban raccoons were infected with several genotypes of hemoplasma similar to *M. haemocanis* (Volokhov et al., 2017); further research should be conducted on the raccoons of Staten Island and the threat they pose to domestic pets and humans.

The low prevalence or absence of infection of other common tick-borne pathogens in hosts was surprising, because other studies found that feral cats, gray squirrels, raccoons, striped skunks, and Virginia opossums were infected with *A. phagocytophilum* (Levin et al., 2002; Keesing et al., 2012), *B. burgdorferi* (only included raccoon and striped skunk, Fish and Daniels, 1990), *B. microti* (Hersh et al., 2012), and *Ehrlichia* spp. (only included raccoons, Dugan et al., 2005). Some studies suggest that small mammals are more effective reservoirs of important human pathogens (i.e. *A. phagocytophilum, B. microti*, and *B. burgdorferi*) compared to medium or large mammals because of different life history characteristics such as body size and reproductive strategy (Ostfeld et al., 2014; Barbour et al., 2015). While opossums were not infected with any microbial pathogens, they were infected with eggs from several nematode species and strongylid macroparasites which could affect host health by making them more susceptible to other pathogens or infections (Ezenwa, 2016).

No tick-borne pathogens were identified from any partially or fully engorged *H. longicornis* larvae removed from infected and uninfected mesomammals. The absence of pathogens in this invasive tick is consistent with a recent metagenomic study that investigated the pathobiome of questing and white-tailed deer-derived *H. longicornis* from the same locations on Staten Island (Tufts et al., in press). The results presented here suggest that while large and medium sized mammals are important hosts for feeding *H. longicornis* ticks, populations of *H. longicornis* in the New York City area have yet to acquire native pathogens present in these mammalian hosts. Furthermore, *H. longicornis* appears unlikely to acquire pathogens through co-feeding with other tick species (*I. scapularis* or *A. americanum*). However, the potential population-level interactions between *H. longicornis* and native tick species are yet to be determined. It is recommended that control efforts to reduce populations of *H. longicornis* in the United States focus on large and medium sized mammals, especially raccoons, opossums, and white-tailed deer.

## Acknowledgements

The authors thank Dr. Peter Donchik, DVM and Matthew Birney for their assistance collecting samples from mesomammals, Anthony DeNicola for his assistance anesthetizing white-tailed deer, and Kevin Zhao for fecal analysis assistance. This publication was supported by the Cooperative Agreement Number U01CK000509, funded by the Centers for Disease Control and Prevention. Its contents are solely the responsibility of the authors and do not necessarily represent the official views of the Centers for Disease Control and Prevention or the Department of Health and Human Services. Columbia University IACUC approved protocols (AABA1469-AABA1472, AAAZ7453, AAAX4454, AAAX4456) and New York State department scientific collector’s permit (LCPSCI-2214).

