## Supplemental Table 1 for "Association of the invasive *Haemaphysalis longicornis* tick with vertebrate hosts, other native tick vectors, and tick-borne pathogens in New York City"

Supplemental Table 1. List of all hosts where *Haemaphysalis longicornis* was found co-feeding with at least one tick of either *Ixodes scapularis, Amblyomma americanum*, or both.

| Date | Site | Host species | *H. longicornis* | | | *I. scapularis* | | | *A. americanum* | | |
|---|---|---|---|---|---|---|---|---|---|---|---|
| | | | A | N | L | A | N | L | A | N | L |
| 27-Aug-19 | LT | Cat | 0 | 0 | 29 | 0 | 1 | 0 | 0 | 0 | 3 |
| 9-Sep-19 | CP | Cat | 0 | 1 | 126 | 0 | 0 | 0 | 0 | 0 | 5 |
| 10-Sep-19 | CP | Cat | 0 | 0 | 16 | 0 | 0 | 1 | 0 | 1 | 7 |
| 10-Sep-19 | CP | Cat | 0 | 2 | 76 | 0 | 0 | 2 | 0 | 0 | 22 |
| 9-Sep-19 | CH | MA | 0 | 0 | 24 | 0 | 0 | 0 | 0 | 2 | 2 |
| 18-Sep-19 | LT | MA | 0 | 0 | 6 | 0 | 8 | 1 | 0 | 3 | 23 |
| 16-Jul-19 | CH | OP | 1 | 1 | 0 | 0 | 0 | 46 | 0 | 0 | 0 |
| 16-Jul-19 | CH | OP | 2 | 19 | 0 | 0 | 1 | 65 | 0 | 0 | 0 |
| 16-Jul-19 | CH | OP | 8 | 19 | 1 | 0 | 3 | 103 | 1 | 0 | 2 |
| 17-Jul-19 | CH | OP | 6 | 317 | 1 | 0 | 4 | 41 | 0 | 0 | 0 |
| 17-Jul-19 | CH | OP | 0 | 10 | 0 | 0 | 1 | 30 | 0 | 0 | 0 |
| 30-Jul-19 | CP | OP | 2 | 0 | 0 | 0 | 0 | 9 | 0 | 0 | 0 |
| 31-Jul-19 | CP | OP | 0 | 3 | 1 | 0 | 3 | 194 | 0 | 2 | 0 |
| 13-Aug-19 | CH | OP | 13 | 42 | 339 | 0 | 0 | 63 | 0 | 0 | 4 |
| 13-Aug-19 | CH | OP | 3 | 31 | 68 | 0 | 1 | 17 | 0 | 1 | 2 |
| 13-Aug-19 | CH | OP | 1 | 7 | 58 | 0 | 1 | 12 | 0 | 0 | 2 |
| 13-Aug-19 | CH | OP | 0 | 0 | 4 | 0 | 0 | 33 | 0 | 0 | 0 |
| 13-Aug-19 | CH | OP | 15 | 43 | 520 | 0 | 4 | 10 | 0 | 1 | 45 |
| 14-Aug-19 | CH | OP | 1 | 1 | 8 | 0 | 0 | 22 | 0 | 0 | 0 |
| 14-Aug-19 | CH | OP | 2 | 2 | 34 | 0 | 0 | 24 | 0 | 0 | 1 |
| 14-Aug-19 | CH | OP | 4 | 36 | 169 | 0 | 0 | 69 | 0 | 1 | 8 |
| 26-Aug-19 | GK | OP | 0 | 0 | 51 | 0 | 7 | 11 | 0 | 0 | 0 |
| 9-Sep-19 | CH | OP | 0 | 0 | 225 | 0 | 0 | 7 | 0 | 0 | 0 |
| 9-Sep-19 | CP | OP | 0 | 1 | 16 | 0 | 0 | 3 | 0 | 0 | 0 |
| 17-Sep-19 | ML | OP | 0 | 2 | 46 | 0 | 0 | 2 | 0 | 0 | 0 |
| 18-Sep-19 | ML | OP | 0 | 0 | 19 | 0 | 0 | 8 | 0 | 0 | 0 |
| 27-Jun-19 | WB | RA | 0 | 1 | 0 | 0 | 7 | 0 | 0 | 0 | 0 |
| 27-Jun-19 | WB | RA | 0 | 1 | 0 | 0 | 2 | 0 | 0 | 1 | 0 |
| 17-Jul-19 | CH | RA | 124 | 24 | 0 | 0 | 0 | 0 | 0 | 2 | 0 |
| 30-Jul-19 | CP | RA | 0 | 6 | 0 | 0 | 0 | 0 | 0 | 5 | 1 |
| 30-Jul-19 | CP | RA | 5 | 7 | 10 | 0 | 1 | 2 | 0 | 14 | 65 |
| 30-Jul-19 | CP | RA | 3 | 2 | 1 | 0 | 1 | 15 | 0 | 11 | 15 |
| 30-Jul-19 | CP | RA | 1 | 3 | 0 | 0 | 0 | 6 | 0 | 6 | 23 |
| 30-Jul-19 | CP | RA | 15 | 15 | 271 | 0 | 1 | 16 | 0 | 26 | 198 |
| 30-Jul-19 | CP | RA | 8 | 9 | 0 | 0 | 0 | 12 | 0 | 12 | 14 |
| 30-Jul-19 | CP | RA | 17 | 7 | 0 | 0 | 0 | 0 | 0 | 8 | 0 |
| 30-Jul-19 | CP | RA | 9 | 13 | 0 | 0 | 0 | 14 | 0 | 24 | 55 |
| 31-Jul-19 | CP | RA | 1 | 2 | 1 | 0 | 0 | 0 | 0 | 5 | 0 |
| 31-Jul-19 | CP | RA | 36 | 13 | 2 | 0 | 0 | 0 | 0 | 17 | 3 |

| | | | | | | | | | | |
|---|---|---|---|---|---|---|---|---|---|---|
| 31-Jul-19 | CP | RA | 12 | 8 | 6 | 0 | 1 | 2 | 0 | 28 | 112 |
| 26-Aug-19 | GK | RA | 0 | 0 | 2 | 0 | 1 | 3 | 0 | 2 | 1 |
| 26-Aug-19 | GK | RA | 0 | 0 | 10 | 0 | 1 | 4 | 0 | 8 | 0 |
| 26-Aug-19 | LT | RA | 0 | 0 | 9 | 0 | 0 | 0 | 1 | 3 | 88 |
| 26-Aug-19 | LT | RA | 1 | 0 | 8 | 0 | 0 | 0 | 0 | 1 | 0 |
| 27-Aug-19 | LT | RA | 0 | 0 | 3 | 0 | 0 | 11 | 0 | 5 | 86 |
| 27-Aug-19 | LT | RA | 0 | 1 | 10 | 0 | 0 | 0 | 0 | 2 | 0 |
| 27-Aug-19 | LT | RA | 0 | 7 | 193 | 0 | 0 | 4 | 0 | 3 | 15 |
| 9-Sep-19 | CP | RA | 0 | 2 | 12 | 0 | 0 | 1 | 0 | 1 | 44 |
| 9-Sep-19 | CP | RA | 0 | 1 | 113 | 0 | 0 | 1 | 0 | 8 | 128 |
| 9-Sep-19 | CP | RA | 0 | 1 | 125 | 0 | 0 | 0 | 0 | 9 | 167 |
| 9-Sep-19 | CP | RA | 0 | 2 | 206 | 0 | 0 | 2 | 0 | 9 | 251 |
| 9-Sep-19 | CP | RA | 0 | 5 | 185 | 0 | 0 | 1 | 0 | 6 | 396 |
| 9-Sep-19 | CP | RA | 0 | 2 | 274 | 0 | 0 | 8 | 0 | 11 | 541 |
| 10-Sep-19 | CP | RA | 0 | 0 | 13 | 0 | 0 | 0 | 0 | 0 | 50 |
| 17-Sep-19 | ML | RA | 0 | 0 | 277 | 0 | 0 | 0 | 0 | 10 | 40 |
| 17-Sep-19 | ML | RA | 0 | 0 | 6 | 0 | 0 | 0 | 0 | 1 | 12 |
| 17-Sep-19 | ML | RA | 0 | 0 | 241 | 0 | 0 | 0 | 0 | 1 | 18 |
| 18-Sep-19 | ML | RA | 0 | 0 | 86 | 0 | 0 | 0 | 0 | 1 | 3 |
| 18-Sep-19 | ML | RA | 0 | 1 | 875 | 0 | 0 | 0 | 0 | 1 | 49 |
| 18-Sep-19 | ML | RA | 0 | 0 | 63 | 0 | 0 | 0 | 0 | 0 | 3 |
| 18-Sep-19 | ML | RA | 0 | 0 | 756 | 0 | 0 | 0 | 0 | 1 | 125 |
| 27-Aug-19 | LT | SK | 0 | 0 | 27 | 0 | 1 | 9 | 1 | 8 | 82 |
| 5-Aug-18 | ML | WD | 71 | 3 | 7 | 0 | 0 | 0 | 5 | 0 | 1 |
| 5-Aug-18 | CSI | WD | 1 | 0 | 29 | 2 | 0 | 36 | 0 | 1 | 1 |
| 5-Aug-18 | CSI | WD | 2 | 0 | 18 | 1 | 2 | 64 | 0 | 0 | 0 |
| 5-Aug-18 | CSI | WD | 2 | 0 | 0 | 1 | 0 | 17 | 2 | 0 | 0 |
| 6-Aug-18 | ML | WD | 19 | 1 | 56 | 0 | 0 | 8 | 0 | 0 | 10 |
| 6-Aug-18 | CSI | WD | 0 | 0 | 28 | 0 | 0 | 21 | 0 | 0 | 0 |
| 6-Aug-18 | CSI | WD | 0 | 0 | 4 | 2 | 1 | 16 | 0 | 0 | 0 |
| 6-Aug-18 | CSI | WD | 1 | 0 | 10 | 1 | 0 | 143 | 1 | 0 | 8 |
| 6-Aug-18 | CSI | WD | 1 | 1 | 93 | 3 | 1 | 95 | 0 | 0 | 1 |
| 6-Aug-18 | CSI | WD | 0 | 0 | 11 | 0 | 0 | 7 | 0 | 0 | 0 |
| 20-Aug-18 | CP | WD | 12 | 41 | 33 | 0 | 0 | 61 | 0 | 7 | 28 |
| 21-Aug-18 | CP | WD | 3 | 9 | 77 | 1 | 2 | 3 | 0 | 4 | 22 |
| 21-Aug-18 | CP | WD | 41 | 13 | 107 | 0 | 0 | 25 | 3 | 1 | 12 |
| 21-Aug-18 | FK | WD | 15 | 6 | 140 | 1 | 0 | 0 | 1 | 0 | 0 |

A = Adult, N = Nymph, L = Larva

Site: CP = Clay Pit Ponds State Park, CH = Conference House Park, ML = Mount Loretto Unique Area, WB = Willowbrook Park, LT = Latourette Park, GK = Great Kills Park, CSI = College of Staten Island, FK = Freshkills Park

Host: GS = Eastern gray squirrel, MA = marmot, OP = opossum, RA = raccoon, SK = striped skunk, WD = white-tailed deer
